# A single dose of antibody-1 drug conjugate cures a stage model of African trypanosomiasis

**DOI:** 10.1101/547208

**Authors:** Paula MacGregor, Andrea L. Gonzalez-Munoz, Fatoumatta Jobe, Martin C. Taylor, Steven Rust, Alan M. Sandercock, Olivia J.S. Macleod, Katrien Van Bocxlaer, Amanda F. Francisco, Francois D’Hooge, Arnaud Tiberghien, Conor S. Barry, Philip Howard, Matthew K. Higgins, Tristan J. Vaughan, Ralph Minter, Mark Carrington

## Abstract

Infections of humans and livestock with African trypanosomes are treated with drugs introduced decades ago that are not always fully effective and often have severe side effects. Here, the trypanosome haptoglobin-haemoglobin receptor (HpHbR) has been exploited as a route of uptake for an antibody-drug conjugate (ADC) that is completely effective against *Trypanosoma brucei* in the standard mouse model of infection. Recombinant human anti-HpHbR monoclonal antibodies were isolated and shown to be internalised in a receptor-dependent manner. Antibodies were conjugated to a pyrrolobenzodiazepine (PBD) toxin and killed *T. brucei in vitro* at picomolar concentrations. A single therapeutic dose (0.25 mg/kg) of a HpHbR antibody-PBD conjugate completely cured a *T. brucei* mouse infection within 2 days with no re-emergence of infection over a subsequent time course of 77 days. These experiments provide a demonstration of how ADCs can be exploited to treat protozoal diseases that desperately require new therapeutics.

**Author Summary:** Here we show that antibody-drug conjugates (ADCs) can be re-purposed from cancer immunotherapeutics to anti-protozoals by changing the specificity of the immunoglobulin to target a trypanosome cell surface receptor. Trypanosomes were used as a model system due to the availability of receptor null cell lines that allowed the unambiguous demonstration that ADCs targeted to a parasite surface receptor could be specifically internalised via receptor-mediated endocytosis. A single low dose of the resulting ADC was able to cure a stage 1 mouse model of trypanosome infection. We have used toxins and conjugation chemistry that are identical to anti cancer ADCs demonstrating the ability to piggy-back onto the huge research efforts and resources that are being invested in the development of such ADCs.

The potential for development of ADCs against a wide range of human pathogens is vast, where only epitope binding sites need vary in order to provide selectivity. This provides a far-reaching opportunity for the rapid development of novel anti-protozoals for the targeted killing of a wide range of pathogens that cause disease worldwide, especially in developing countries.

## Introduction

Infection with African trypanosomes causes disease in humans, livestock and wild animals. At least seven species are able to infect livestock but only *Trypanosoma brucei* subspecies normally infect humans: *T. b. gambiense* and *T. b. rhodesiense* cause chronic or acute Human African Trypanosomiasis (HAT) respectively (1). New drug treatments are required for human treatment, the drugs currently used require multiple administrations over periods of weeks and all can have severe side effects (reviewed in (2–4)).

Without intervention, infection persists as the trypanosomes have evolved a population survival strategy based on antigenic variation of the variant surface glycoprotein (VSG) that is present as a densely packed coat on the external face of the plasma membrane. Receptors for host nutrient macromolecules are integrated in the VSG coat, such as the HpHbR which is involved in haem acquisition through binding and subsequent endocytosis of host haptoglobin-haemoglobin(5). Primate-specific innate immune protein complexes have evolved to exploit this nutrient uptake and kill most isolates of *T. brucei* (5). The two complexes, Trypanolytic Factor 1 and 2 (TLF1 and TLF2), each contain two primate-specific proteins, apolipoprotein L1 (apoL-1) (6) and haptoglobin-related protein bound to haemoglobin (HprHb) which acts as a molecular mimic of HpHb(7–10). HpHbR binds and internalises TLF1 and the toxin apoL-1 kills the trypanosome (5, 11). Human infective trypanosomes have evolved counter-measures to the TLFs(12–19).

The binding of a host macromolecule to a receptor, followed by the internalisation of the complex, provides a potential route to specifically deliver therapeutics into trypanosome cells. Entry of TLF1 via the HpHbR and the release of a cytotoxin after internalisation is analogous to the mode of action of ADCs (20), a growing class of therapeutics, particularly used in applications in oncology(21–23) and also with demonstrated potential as anti-bacterials(24, 25). An early attempt to develop ADCs against the intracellular American trypanosome, *Trypanosoma cruzi*, used chlorambucil conjugated to polyclonal IgGs purified from chronically infected rabbits (26) and, while results were promising, this was only partially successful. More recently, antibody therapeutics against African trypanosomes based on single domain antibodies derived from camelid immunoglobulins (nanobodies) recognising some, but not all, VSGs (27, 28) have also been developed. One study used a nanobody apoL-1 fusion protein that was curative in mouse infections(29). In another two studies, nanobodies were used to create nanoparticles containing pentamidine, one of the current drugs used to treat trypanosome infection. These particles bound VSG and were successfully taken up into the endocytic pathway, the concentration required for cure was 10 to 100-fold lower than free pentamidine over a course of four doses (30, 31). However, the variability of the VSG molecules and underpinning antigenic variation will almost certainly limit their effectiveness as targets for therapeutic delivery.

Here we have developed a recombinant human anti-trypanosome-HpHbR antibody conjugated to a PBD toxin, selected so that recognition of the trypanosome would be independent of the VSG identity. This approach also strategically exploits advances in anti-cancer ADC development. The antibody-PBD conjugate was effective at killing trypanosomes in culture at picomolar concentrations whereas killing of human cell lines required more than 100,000-fold higher concentrations. A single low dose (0.25 mg/kg) of one of the ADCs resulted in a long-term cure in the standard mouse model of trypanosome infection(32, 33) with no apparent adverse effects.

## Results

HpHbR was chosen as a target for ADCs for two reasons: first it is responsible for receptor mediated endocytosis of ligands larger than IgGs and structural information suggested it is accessible to external antibodies (34, 35); second, a cell line with both HpHbR alleles deleted (HpHbR −/−) was available as a control for specificity. HpHbR −/− cell lines have little or no growth phenotype in culture (5, 34), although they are attenuated in the murine experimental model of infection (5).

### Identification of single chain variable fragments recognising the N-terminal domain of the haptoglobin-haemoglobin receptor

In *T. brucei*, the mature HpHbR has a large N-terminal domain (264 residues) that contains the HpHb binding site (34) and a small C-terminal domain (79 residues) attached to the plasma membrane by a glycosylphosphatidylinositol anchor. Recombinant HpHbR N-terminal domain (34) was used for phage display affinity selection from a single chain variable fragment (scFv) library. Specificity for HpHbR was confirmed using phage ELISA and sixteen distinct scFvs were identified (Figure 1A).

**Figure 1:**
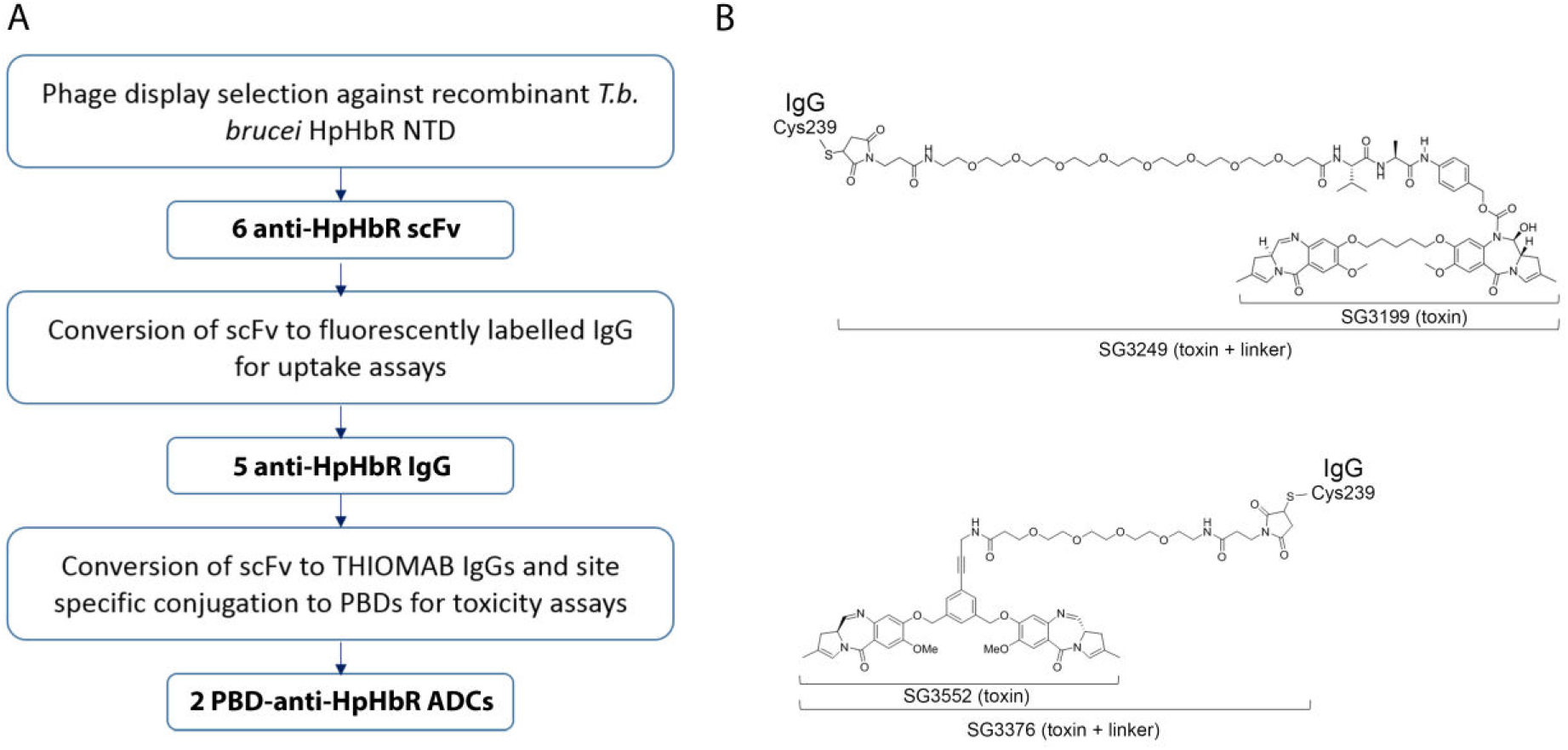
The generation of ADCs that target the *T. brucei* HpHbR. (A) Workflow for the generation of anti-trypanosomal ADCs. (B) Structures of the two PBD toxins (SG3199 and SG3552) and their corresponding toxins plus linker derivatives (SG3249 and SG3376) used in this study. Note that the linker of SG3249 contains a cleavable dipeptide motif whereas the linker of SG3376 does not.

### HpHbR antibodies are internalised by receptor mediated endocytosis

Six of the scFvs (S1 Figure) were reformatted as human IgG1 for further analysis. To determine whether any of these IgGs were endocytosed by trypanosomes in a receptor dependent manner, each was labelled with Alexa fluor-594 and incubated with either *Trypanosoma brucei*, Lister 427, HpHbR wild-type or HpHbR −/− cells in culture for 2 hours in the presence of the lysosomal protease inhibitor FMK-024. A control IgG1 with an unrelated specificity (NIP228) was used in parallel.

Internalisation was monitored by microscopy (Figure 2) and at 10 nM IgG1 five of the six HpHbR antibodies were endocytosed by wild-type cells but not by HpHbR−/− cells and localised to a compartment consistent with the lysosome. There was no internalisation of the control antibody in either cell line at 10 nM. Hence, five of the antibodies were internalised by receptor mediated endocytosis demonstrating that they recognised epitopes on HpHbR that are accessible on live cells. The sixth HpHbR antibody (Tb086) showed limited internalisation and was not used further.

**Figure 2:**
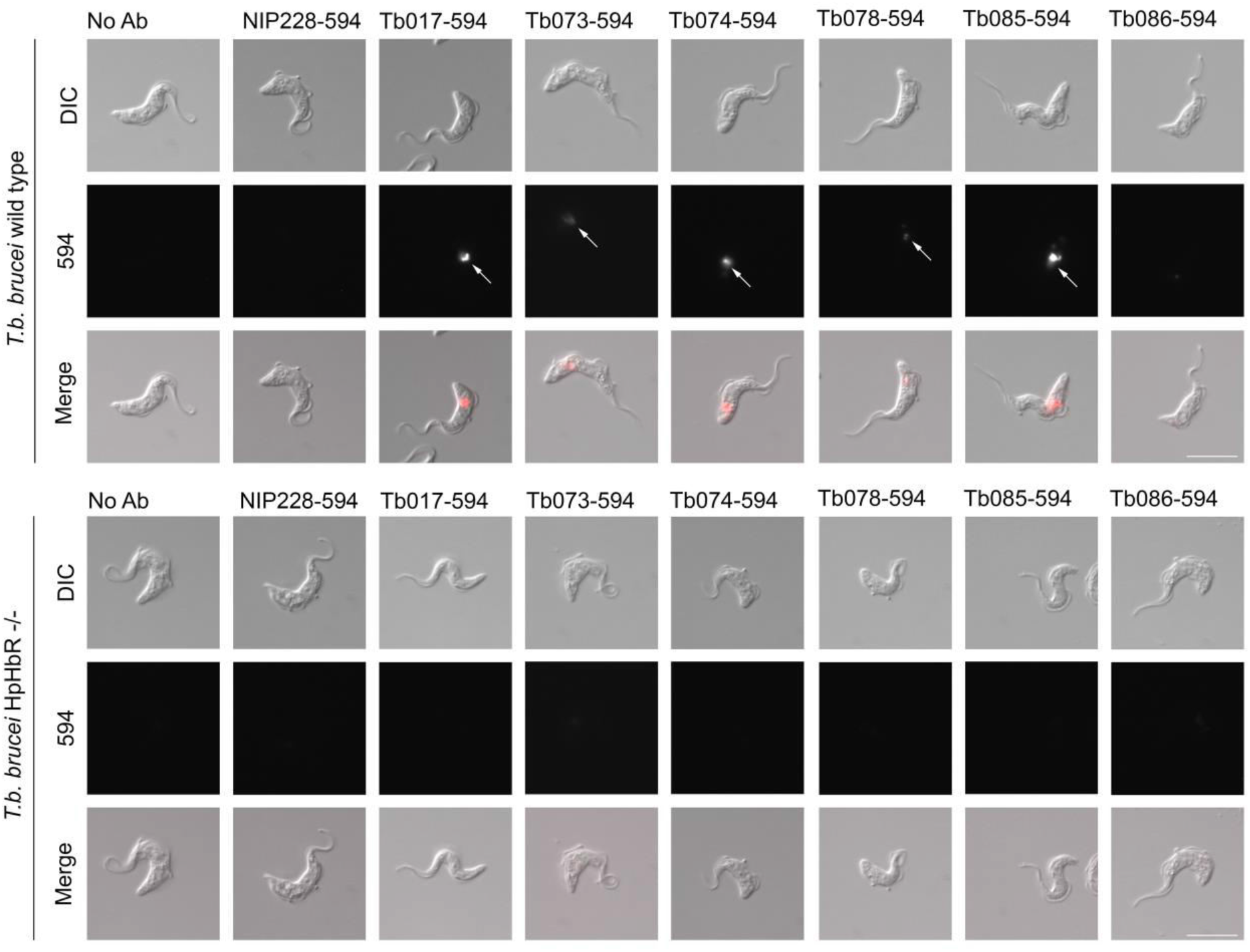
Receptor mediated endocytosis of humanised anti-HpHbR IgG1s. Uptake of Alexa594-labelled antibodies into *T. b. brucei* Lister 427 *HpHbR* wild type and −/− cells was monitored by microscopy. Uptake of five of the seven selected antibodies was detected at 10 nM in wild-type (indicated by arrows in upper panel) but not in HpHbR −/− cells (lower panel). No specific uptake of the remaining antibody (Tb086) or a control antibody (NIP228) was detected. Scale bar represents 10 μm.

### Toxin-conjugated HpHbR-targeting antibodies kill trypanosomes at picomolar concentrations

The receptor-mediated endocytosis of these HpHbR antibodies was then exploited to assess the effectiveness of ADCs against *T. brucei in vitro.* Two PBDs, SG3199 and SG3552 (ref(36)) (Figure 1B), were used in these experiments; each was used as a toxin-linker derivative, SG3249 and SG3376 respectively (Figure 1B), for antibody conjugation. PBDs are DNA minor groove binding toxins (37–40) and were chosen as trypanosomes have a highly complex mitochondrial genome formed from a network of thousands of concatenated DNA circles and are consequently susceptible to DNA binding toxins. This sensitivity is illustrated by the original patent on ethidium bromide as a treatment for trypanosome infection and ethidium derivatives are still used for animal trypanosomiasis (41, 42).

To assay for trypanocidal activity, cultures of *T. brucei* were incubated with a range of concentrations of the anti-HpHbR-PBD conjugates over 48 hours. Growth was measured as percentage proliferation compared to no treatment, with 0% relative to controls representing no viable cells observed, and IC_50_ values calculated.

Initial experiments were designed to identify the most effective HpHbR antibody and used the PBD, SG3199. Free SG3199 had an IC_50_ of ~1 pM (Figure 3A, Table S1), this confirmed its toxicity towards trypanosomes and indicated that it is freely cell permeable. Prior to conjugation to the IgGs, SG3199 was modified by the addition of a linker to facilitate conjugation and release in the lysosome after proteolysis to produce SG3249(43) (Figure 1B). Free SG3249 had an IC_50_ of ~240 pM (Figure 3A, Table S1); presumably the hydrophilic nature of the linker meant that cell access via passive diffusion was reduced. Antibody-SG3249 conjugates were prepared for the five HpHbR antibodies selected in the uptake experiment above and the NIP228 IgG control, following IgG engineering to contain a surface exposed cysteine residue at position 239 in the heavy chain CH2 domain for conjugation to PBD molecules(44) (Figure 1B). The HpHbR antibody-SG3249 conjugates all killed trypanosomes with IC_50_ values between 9 and 86 pM compared to 2100 pM for the control NIP228-SG3249 conjugate (Figure 3A and Table S1), demonstrating targeted cell killing by HpHbR antibody-PBD conjugates. The two most potent antibodies were Tb074 and Tb085 with IC_50_ values of 17 and 9 pM respectively and they were selected for further experiments.

**Figure 3:**
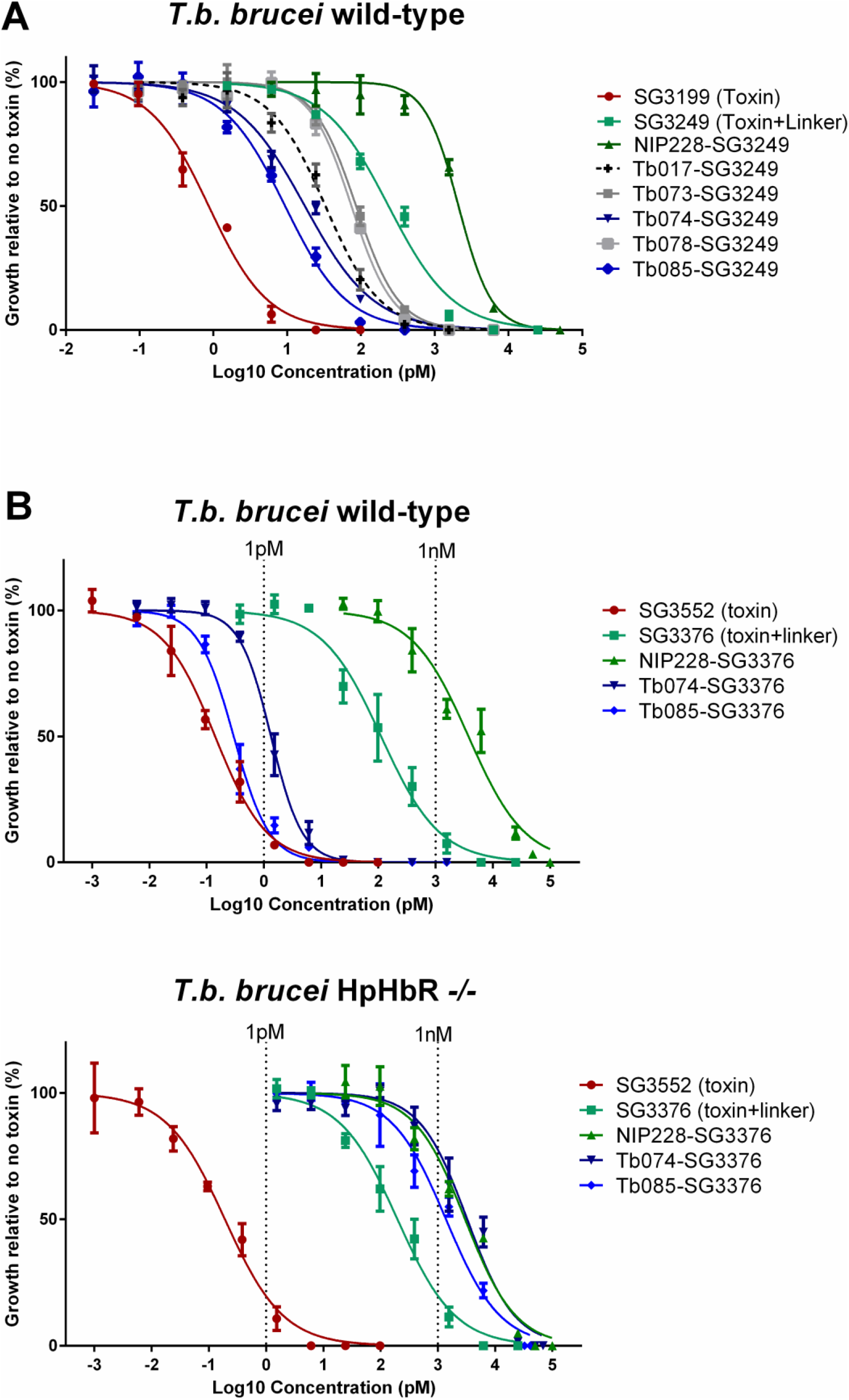
HpHbR antibody-PBD conjugates result in *T. brucei* cell death at low picomolar concentrations *in vitro* in a HpHbR-dependent manner. (A) Toxin SG3199 kills *T. b. brucei* wild type cells at sub-picomolar concentrations (IC_50_ 0.86 pM), killing activity is reduced by the addition of a linker (SG3249 IC_50_ 236.0 pM). Conjugation of SG3249 to a non-specific control antibody (NIP228) further reduces trypanosome killing activity to low nanomolar concentrations (IC_50_ 2.1 nM) whereas conjugation of SG3249 to antibodies that target the HpHbR increased killing activity to low picomolar concentrations (IC_50_ values range from 86 pM for Tb073-SG3249 to 9.4 pM for Tb085-SG3249). All assays were carried out in triplicate over 48 hours. Lines represents nonlinear regression lines of best fit on Log-10 transformed data. Error bars represent standard error of the mean (s.e.m.), n=3 biological replicates (carried out in parallel). (B) Toxin SG3552 kills *T. b. brucei* wild type and HpHbR −/− cells with sub-picomolar IC_50_ concentrations The IC_50_ is increased by orders of magnitude by the addition of a linker (Table 1). Conjugation of SG3376 to a non-specific control antibody (NIP228) further increases the IC_50_ to nanomolar concentrations in both trypanosome cell lines. HpHbR antibody SG3376 conjugates have an IC_50_ in the low/sub picomolar range for wild type *T. b. brucei.* In contrast, IC_50_ values with *T. b. brucei* HpHbR −/− cells remained similar to the control ADC. All assays were carried out in triplicate over 48 hours. Lines represents nonlinear regression lines of best fit on Log_10_ transformed data. See Table 1 for corresponding IC_50_ values. Error bars represent s.e.m., n=3 biological replicates (carried out in parallel).

The next set of experiments used PBD SG3552 and its linker-derivative SG3376 (45, 46) (Figure 1B). This toxin-linker combination was chosen as it was designed to have fewer off-target effects (45, 47) and was shown to be more potent against trypanosomes in preliminary experiments. Three antibody-SG3376 conjugates were prepared from Tb074, Tb085 and NIP228 and all were tested for trypanocidal activity as above but using HpHbR wild type and −/− cell lines (Figure 3B and Table 1). SG3552 killed trypanosomes with IC_50_ values of 0.14 pM in wild type and 0.2 pM in HpHbR −/− cell lines; the addition of the linker to make SG3376 reduced the toxicity to 112 pM and 197 pM in wild type and −/− cell lines respectively, again presumably due to the increase in hydrophilicity conferred by the linker reducing passive cell entry. The antibody conjugates Tb085-SG3376 and Tb074-SG3376 were effective in killing wild-type trypanosomes with IC_50_ values of 0.3 pM and 1.3 pM respectively. In contrast both were far less effective against HpHbR −/− cells with IC_50_ values of 1390 pM and 3270 pM showing that the action of the ADC is dependent on HpHbR expression. The action of the NIP228-SG3376 conjugate was unaffected by HpHbR expression and had an IC_50_ of 3750 pM and 3000 pM in HpHbR wild type and -/cells respectively. Taken together these findings showed that HpHbR antibody-SG3376 conjugates are highly effective in killing trypanosomes through a mechanism whereby the presence of the receptor increases specificity by several thousand-fold over the action of non-specific antibody-SG3376 conjugates.

**Table 1:** IC_50_ values (pM) of SG3552-based toxins and ADCs against *T.brucei* cell lines and a human Jurkat cell line. The IC_50_ values of toxin SG3552, toxin plus linker SG3376, a control ADC (NIP228-SG3376) and two anti-trypanosome ADCs targeting the *T. b. brucei* HpHbR (Tb074-SG3376 and Tb085-SG3376) against *T. b brucei* wild type and *T. b brucei* HpHb −/− (Figure 3B) were calculated. Values in bold are best-fit IC_50_ values, the range is the 95% confidence intervals. It was not possible to calculate accurate IC_50_ values for the Jurkat cell line due to lack of saturation of the cell killing assay and so all were conservatively estimated as greater than 50 nM from the data in S2 Figure. All values are shown to 3 significant figures.

| Cell line | IC<sub>50</sub> (pM) |  |  |
| --- | --- | --- | --- |
|  | *T. b. brucei* | *T. b. brucei*<br>*HpHbR -/-* | Human Jurkat |
| **SG3552 Toxin** | **0.14**<br>(0.11-0.18) | **0.195**<br>(0.14-0.27) | **19.6**<br>(10.8-35.8) |
| **SG3376 Toxin plus linker** | **112**<br>(76.7-163) | **197**<br>(145-268) | **>50,000** |
| **NIP228-SG3376 Control** | **3750**<br>(2610-5380) | **3000**<br>(2260-3960) | **>50,000** |
| **Tb074-SG3376** | **1.33**<br>(1.16-1.53) | **3270**<br>(2400-4450) | **>50,000** |
| **Tb085-SG3376** | **0.297**<br>(0.25-0.36) | **1390**<br>(1030-1880) | **>50,000** |

To assess whether the HpHbR antibody-SG3376 conjugates have specificity for trypanosomes over mammalian cells in culture, PBD toxin SG3552 and antibody-SG3376 conjugates were assessed for toxicity against a range of human cell lines. SG3552 was toxic to all cell lines assayed at picomolar concentrations (S3 Figure), the most sensitive was the Jurkat cell lines with an IC_50_ value of 19.6 pM, around 100-fold less-sensitive than the *T. brucei* cell lines (Table 1). This was expected: trypanosomes are particularly sensitive to many DNA damaging toxins as described above. The NIP228-SG3376, Tb074-SG3376 and Tb085-SG3376 conjugates all had IC_50_ values that were conservatively estimated to be >50 000 pM (S3 Figure). The IC_50_ values of the two HpHbR antibody-SG3376 conjugates for the human cell lines was at least 50,000 times greater than those for trypanosomes (Table 1).

### A single Tb085-SG3376 administration results in the clearance of trypanosome infection in mice

Based on the specificity and potency observed in the above experiments, Tb085-SG3376 conjugate was chosen to determine anti-HpHbR-toxin conjugate efficacy in a mouse model of *T. b. brucei* infection. Mice were infected with a pleomorphic trypanosome cell line, *T. b. brucei* GVR35-VSL2, that expresses a luciferase transgene (PpyRE9h) to facilitate measurement of infection in live animals over a prolonged time course using bioluminescence imaging (BLI) (32, 33). This method has the advantage that it detects trypanosomes in the bloodstream and tissues. Fifteen mice were infected with trypanosomes and imaged on day 3 post infection to provide a pre-treatment BLI signal level indicative of the whole-body infection burden measured as photons per second (p/s) after administration of luciferase substrate.

All infected mice had a total flux of between 2.5×10^9^ and 5.9×10^9^ p/s with the exception of a single mouse which had a lower level of infection at 3×10^7^ p/s. Subsequent to imaging, on day 3, groups of five mice were then treated with (1) 0.25 mg/kg Tb085-SG3376 or (2) 0.25 mg/kg NIP228-SG3376 or (3) PBS alone. Three uninfected mice were used as negative controls for the BLI.

Infection levels were assessed by BLI on days 4, 5, 6 and 7, and then at regular further time points (Figure 4, S4 Figure, S5 Figure). Within the first day posttreatment the BLI signal in Tb085-SG3376-treated mice had dropped 3-fold relative to the pre-treatment signal whilst control mice (NIP228-SG3376 or PBS alone) had increased more than 2-fold. These control mice remained infected with a BLI signal consistent with a first and second wave of parasitaemia, characteristic of trypanosome infection dynamics (48, 49). At day 14 (11 days post-treatment), control mice were culled at a humane endpoint, as the BLI signal represented a parasite burden that would invariably lead to clinical symptoms of trypanosomiasis and death (33).

**Figure 4:**
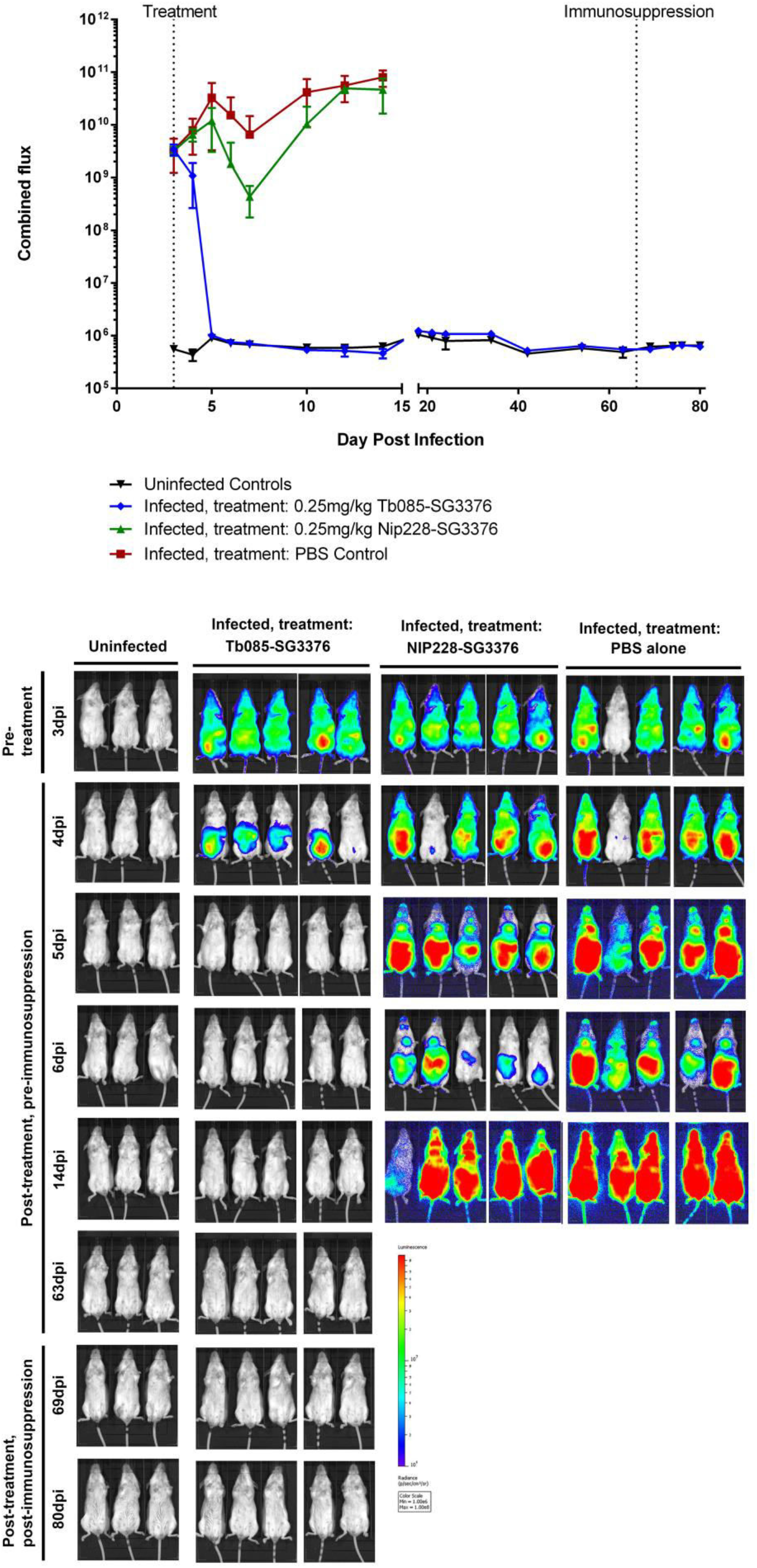
A single low dose of Tb085-SG3376 was able to cure infection in a mouse model of trypanosomiasis. Three groups of 5 mice were infected with pleomorphic *T. b. brucei* GVR35-VSL2 cells (32, 33), which allow for parasite burden in live mice to be assessed over a time course by bioluminescent imaging (BLI). BLI was performed prior to any treatment at 3 dpi and then at regular time points following treatment on 3 dpi with a single intravenous dose of (1) 0.25 mg/kg Tb085-SG3376 (n=5), (2) 0.25 mg/kg NIP228-SG3376 (n=5) or (3) PBS alone (n=5). Unlike the control-treated mice, Tb085-SG3376 treatment caused a decrease in the luminescent signal to that obtained from uninfected control animals within 2 days and this remained the case for the duration of the infection, including following the immunosuppression of Tb085-SG3376 treated mice at 66 dpi. Mice treated with NIP228-SG3376 or PBS were culled at a humane endpoint on day 14. (A) Quantification shown is the combined (dorsal + ventral) luminescence over the whole mouse in photos per second (p/s). The corresponding quantification data from the 18 individual mice are shown in S5 Figure. Error bars represent standard deviation. Downward error bars are missing from 4 data points due to scale constraints. (B) For each group of mice selected ventral images for the BLI are shown. Corresponding dorsal images of the same mice are shown in S4 Figure. The scale bar represents the photons emitted at any given point on the image. Exposure times range from 0.5 seconds (for heavily burdened mice) to 5 minutes (for uninfected animals). One mouse in the PBS control group had a lower BLI signal than all other infected mice at 3 dpi (S5 Figure). In the image shown here this mouse appears negative, however, this is due to the low exposure time required for adjacent mice.

In contrast, the BLI signal in mice in group 1 (treated with Tb085-SG3376) had decreased to the level of uninfected controls by 2 days post-treatment. The BLI signal remained indistinguishable from the uninfected controls for 60 days posttreatment and the mice continued to appear healthy throughout the experiment, not showing any external symptoms of clinical trypanosomiasis. To determine if Tb085-SG3376 treated mice were harbouring very small numbers of trypanosomes that were kept in check by the mouse adaptive immune response, the mice were immunosuppressed with a single dose of cyclophosphamide on day 66 post-infection and BLI measurements made on days 69, 74, 76 and 80 post-infection; no trypanosomes were detected (Figure 4, S4 Figure, S5 Figure). On day 80 postinfection mice were culled and BLI was performed on mouse tissues post-necropsy; again no trypanosomes were detected in any tissue (S6 Figure). Finally, both a blood sample and a section of brain tissue from each of the five mice treated with Tb085-SG3376 were incubated in trypanosome culture medium for one month; in no case were any trypanosomes then detected. Together, these observations and measurements indicate that a single dose of Tb085-SG3376 was sufficient to cure infection in 5/5 mice in the experimental group.

## Discussion

African trypanosomes proliferate in the bloodstream and tissue spaces of their mammalian hosts where they are continually exposed to the adaptive immune response. The trypanosome cell surface is covered by a densely packed coat of VSG that underpins persistence of infection by antigenic variation. The VSG coat must be permissive for receptor mediated endocytosis of host macromolecules as nutrients and here this has been exploited for the delivery of an ADC. The HpHbR was chosen for this study as: (i) it is a natural route for uptake of the trypanolytic factors(5), which kill sensitive trypanosomes strains in human serum; (ii) it is accessible to ligands larger than IgG (5); (iii) it has a known structure (34, 35); (iv) HpHbR null cell lines grow at a normal rate in culture (5, 34) and were an ideal control for specificity of uptake. We found that HpHbR monoclonal antibodies are taken up into HpHbR wild type cells but not HpHbR −/−cells, proving that receptors for host macromolecules are accessible on live trypanosomes. These same antibodies conjugated to a PBD were able to kill trypanosomes in culture at pM concentration in a manner that was dependent on HpHbR expression. Significantly higher doses, were needed to kill a panel of mammalian cell lines. Finally, in the mouse model of infection, a single administration of an anti-HpHbR ADC was sufficient to cure the infection.

The findings here have validated an approach that builds on the considerable progress in anti-cancer ADCs and repurposing into an anti-protozoal simply involves the development of pathogen specific antibodies. The use of ADCs here was specifically based on those developed in oncology. Currently, ADCs are used in the clinic against Hodgkin lymphoma (Brentuximab vedotin) (22) and HER2-positive breast cancer (ado-trastuzumab-emtansine) (50). Many others are in pre-clinical development or clinical trials, including ADCs against a range of cancers that incorporate PBDs, including SG3249, one of the toxins used in this study (51–53).

The success of the experiments above lead to the question of whether this is a realistic approach for development of therapeutics for trypanosome and other protozoan infections. Amongst the key challenges in generating ADCs for applications in oncology is ensuring minimal off-target toxicity and so, as well as through ADC chemistry, low doses are desirable (reviewed in (54)). The single dose of 0.25 mg/kg was selected in these experiments as a proof-of-concept because it is at the lower end of effective oncological ADC treatment in mice(55) and is well below the anticipated maximum tolerated dose (56). The minimum efficacious dose achievable with the anti-HpHbR ADC was not tested in this study and it is likely that the targeting of parasites will be achieved using lower doses than required for oncology for two key reasons. First, in contrast to the surface of cancer cells, parasite-specific surface receptors are entirely different from host cell surface receptors leading to highly selective uptake of the antibody into the pathogen. Second, the effectiveness of the ADC in this study was enhanced by the sensitivity of trypanosomes to DNA-binding agents, in comparison to host cells. Together these led to a 100,000-fold difference in toxicity between trypanosome and human cells *in vitro.* These considerations will also apply to other protozoal pathogens providing a suitable target can be identified.

Disease caused by *T. brucei* infection has two stages: in stage 1 trypanosomes are excluded from the central nervous system (CNS) by the blood brain barrier (BBB) while in stage 2 infections trypanosomes enter the CNS. In the experimental model used here, we have tested the ability to clear a stage 1 infection. Would ADCs be able to target trypanosomes in the CNS? While administered intravenous antibodies are present in the CNS at less than 0.1% of the concentration in the blood in murine models (57, 58) increased BBB permeability has been observed in murine models of neurological-stage trypanosomiasis (59–61), which will increase the CNS concentration of administered antibodies. Further, bifunctional fusion antibodies that can cross the blood-brain barrier have been reported (57).

It is worth contrasting a potential ADC treatment with the current effective drug regimens for trypanosomiasis. Pentamidine, the current stage 1 *T. b. gambiense* treatment, is administered to patients intramuscularly at 4 mg/kg over 7 days, although it has been shown to clear a mouse model of *T. b. brucei* infection at 2.5 mg/kg over four intraperitoneal injections (30, 62). For stage 2 *T. b. gambiense* infection, the current nifurtimox eflorithine combination therapy involves oral nifurtimox 15 mg/kg/day for 10 days plus eflornithine infusions 400 mg/kg/day for 7 days (for a 50 kg adult this is 20 g eflornithine per day) (63). A single dose of ADC would clearly be an improvement.

Considerable resources are being used for the optimisation, assessment and clinical trials of oncology ADCs. It is difficult to imagine such resources being available for the developmental pipeline of therapeutics against protozoal pathogens that primarily affect developing countries. Both cancer and protozoal pathogens are eukaryotic cells and so the oncology-based strategies that take advantage of the cell biology of cancer cells are often applicable to protozoa. Therefore, the scope for benefiting from oncology developments is clear, particularly where the drug (such as PBDs, as used in this study) do not deviate from oncology ADCs that are under development. If simply modifying the epitope binding site can allow anti-cancer ADCs to be repurposed then they could realistically be developed as a novel class of therapeutics for protozoan pathogens. The cell surfaces of protozoan pathogens are often particularly well studied due to the biological interest in their role in host:parasite interactions and therefore the literature contains a reservoir of potential targets (for example (64–68)). It is also worth noting that the production cost of ADCs is far less than often realised (69–73).

In summary, we have demonstrated that a single dose of an ADC, shown to specifically operate through the HpHbR was able to completely cure an infection in a stage 1 trypanosomiasis model. These type of agents have the potential for development for use to treat trypanosome infection in humans, and in the longer term livestock animals. Furthermore, this work illustrates that developments in oncology ADCs can be applied to protozoal pathogens, the causal agents of many neglected diseases in need of new therapeutics.

## Materials and Methods

### Phage display selection of anti-HpHbR N-terminal domain single chain variable fragments

Recombinant HpHbR N-terminal domain (NTD) was expressed as previously described (34) and a scFv antibody library was used to perform soluble and panning phage display selections (74). Briefly, panning selections were performed by coating 5 μg/mL biotinylated HpHbR NTD on to a single well of a streptavidin-coated 96-well plate or 10 μg/mL non-biotinylated HpHbR NTD on to a single well of a Nunc Maxisorp plate overnight at 4°C. Coated wells were washed three times with phosphate buffered saline (PBS) prior to incubation for 1hr at room temperature with 3% Marvel skimmed milk powder in PBS. Next, 1 × 10^12^ phage particles in 6% Marvel in PBS were added to each coated well and incubated for 1 hr at room temperature. The wells were washed five times with PBS containing 0.1% Tween-20 and five times with PBS prior to elution and recovery of phage. For soluble selection, phage were pre-incubated with magnetic beads in 3% Marvel in PBS at room temperature for 1 hour. Subsequently, the magnetic beads were removed and the phage-containing supernatant was incubated with biotinylated HpHbR NTD at room temperature for 1 hour. Streptavidin magnetic beads were subsequently added to the reaction and incubated at room temperature for 5 minutes. The magnetic beads were washed five times with 0.1% Tween-20 in PBS. For all selections, phage were eluted with 10 μg/ml trypsin in PBS for 30 minutes at 37°C. Exponentially grown TG1 *E.coli* cells were infected with the eluted phage and grown overnight at 30°C on agar plates containing ampicillin. *E. coli* colonies were harvested from the bioassay plates and phage particles were rescued by super-infecting with M13 KO7 helper phage and used in the next round of selection. In total, three serial rounds of selection were performed.

### Phage ELISA

Individual phage were produced from *E. coli* and assayed, by phage ELISA, against TbHpHbR NTD in parallel with BSA and streptavidin. Briefly, 10 μg/ml of each protein was coated onto Nunc Maxisorp plates and 5μg/mL of each biotinylated protein was coated onto streptavidin-coated plates overnight at 4°C. Plates were washed three times with PBS before being incubated with 3% Marvel in PBS for 1 hour at room temperature. Phage containing supernatants were blocked with an equal volume of 6% Marvel in 2xPBS for 1 hour at room temperature. Coated plates were washed three times with PBS and incubated with 50 μl of blocked phage supernatants for 1hr at room temperature. Plates were washed three times with 0.1% Tween 20 in PBS and bound phage were detected using an anti-M13 horseradish peroxidase conjugated antibody and colorimetric substrate. Rabbit polyclonal anti-TbHpHbR antibody was used as a positive control and detected with mouse anti-rabbit IgG HRP.

### Generation of full length human IgG1 and THIOMABS

Selected scFvs were converted to full length human IgG1s using standard molecular biology techniques. Plasmids encoding secreted antibody (75) were purified by protein A affinity chromatography. Recombinant antibody was labelled with Alexa Fluor 594 following the manufacturer’s instructions (Life technologies). Standard molecular biology techniques were used to introduce a cysteine residue at position 239 in the CH2 domain of each heavy chain (44). Recombinant THIOMABs were expressed and purified as detailed for full length IgG1.

### PBD conjugation to THIOMABs

The HpHbR THIOMABS and a NIP228 negative control were reduced by the addition of a forty fold molar excess of tris(2-carboxyethyl)phosphine (TCEP) in PBS, 1 mM EDTA, pH 7.2 for 4 h at 37°C. TCEP was subsequently removed and the THIOMABS were re-oxidised with a twenty times molar excess of dehydroascorbic acid for 4h at 25°C. A ten times molar excess of toxin plus linker was added and incubated for 1 h at 25 °C, the reactions were quenched by the addition of excess of N-acetyl-L-cysteine. The resultant ADCs were formulated in PBS, pH 7.2 after ultrafiltration to removed excess toxin. ADCs were characterized by determination of monomeric purity by size exclusion chromatography (Table S2), drug-antibody-ratio (DAR) by RP-HPLC chromatography (Table S2) and molecular mass (by LC-MS of the reduced ADCs) (S5 Figure)

### Trypanosome cell culture

*T. b. brucei* Lister 427 bloodstream cells were grown in HMI-9 salts plus 10% foetal calf serum (FCS) at 37°C with 5% CO_2_ (76). The *T. b. brucei* Lister 427 HpHbR −/− cell line used here has been described previously (34).

### Internalisation of fluorescently labelled IgGs into live cells

For *T. b. brucei* uptake assays 1 × 10^6^ cells per assay were incubated with 10 nM Alexa Fluor 594-labelled IgG in 300μl HMI-9, 10% FCS, 2μM FMK-024 protease inhibitor for 1.5 hours at 37°C. Cells were washed once in HMI-9, 10% FCS then fixed in 1% PFA for 10 minutes at room temperature and resuspended in PBS. Internalisation was determined by microscopy using a Zeiss Imager M1 microscope and analysed with AxioVision Rel 4.8 software.

### In vitro *trypanosome cell-killing assays*

*T. b. brucei* Lister 427 wild-type or HpHbR −/− cell lines were incubated at 1 × 10^4^ cells/ml in triplicate with PBDs or ADCs for 48 hours before cells were counted and growth was calculated relative to an untreated control for each cell line. All assays contained 0.5% DMSO. Data were Log_10_ transformed and nonlinear regression lines of best fit and IC_50_ values were calculated using GraphPad Prism 6.

### CellTiter-Glo Luminescent Cell Viability Assay

*In vitro* viability cell assays were performed with primary and transformed human cell lines: Raji (ECACC), Jurkat E6.1 (ATCC), NHLF (LONZA) and HUVEC (LONZA). These cell lines were mycoplasma tested and authenticated by PCR using human 16-marker short tandem repeat profiling and interspecies contamination test by IDEXX (Columbia, MO). Cells seeded at 2 × 10^5^ cell/ml (Raji and Jurkat) and at 2 x 10^3^ cell/ml (NHLF and HUVEC) in 96 well plates were incubated with the SG3552 toxin, the toxin+linker SG3376 and the corresponding ADCs (Tb074-SG3376, Tb085-SG3376 and NIP228-SG3376). All assays contained 0.5% DMSO. After 96 hours, the number of viable cells in culture was measured using the CellTiter-Glo 2.0 luminescent cell viability assay and read in Envision plate reader. Growth was calculated relative to an untreated control for each cell line. Data were Log_10_ transformed and nonlinear regression lines of best fit and IC_50_ values were calculated, where possible using GraphPad Prism 6.

### Mouse infection and bioluminescent imaging of trypanosome infection

Pleomorphic *T. b. brucei* GVR35-VSL2 bloodstream forms were cultured and maintained at 37°C/5%CO_2_ in HMI-9 medium supplemented with 20% FBS, 1μg/ml puromycin and 1% methyl cellulose (33). Parasites were maintained at <1 × 10^6^ ml^−1^ and were not cultured for more than three passages prior to mouse infection.

Mice were purchased from Charles River (UK). They were maintained under specific pathogen-free conditions in individually ventilated cages with a 12 hour light/dark cycle and access to food and water *ad libitum.* Female BALB/c mice aged 8 to 12 weeks were infected intraperitoneally with 3×10^4^ *T. b. brucei* GVR35-VSL2 cells (33). Three groups of five mice were infected. On day 3 post infection the mice were imaged to obtain the pre-treatment infection level. Five mice received 0.25 mg/kg Tb085-SG3376, five mice received PBS alone and five mice received 0.25 mg/kg NIP288, all intravenously. A group of three mice was not infected.

Imaging was carried out by intraperitoneal injection of 150 mg/kg D-luciferin. After 5 minutes, mice were anaesthetised with 2.5% (v/v) gaseous isofluorane in oxygen. The mice were transferred to the IVIS Illumina and imaged using LivingImage 4.3. software (PerkinElmer). Exposure times were determined automatically and varied between 0.5 s and 5 min depending on the radiance. After imaging, mice were allowed to recover and transferred back to their cages.

At 66 days post-infection, Tb085-SG3376 treated mice were immunosuppressed with a single intraperitoneal dose of cyclophosphamide (200 mg/kg).

### Ethics statement

All animal work was performed under UK Home Office licence 70/8207 and approved by the London School of Hygiene and Tropical Medicine Animal Welfare and Ethical Review Board. All protocols and procedures were conducted in accordance with the UK Animals (Scientific Procedures) Act 1986.

## Acknowledgments

This work was supported by Medical Research Council Project Grant MR/L008246 to MC and MH. PM is a BBSRC David Phillips Fellow (BB/P010849/1).

## Supporting information Figure Legends

**S1 Figure:**
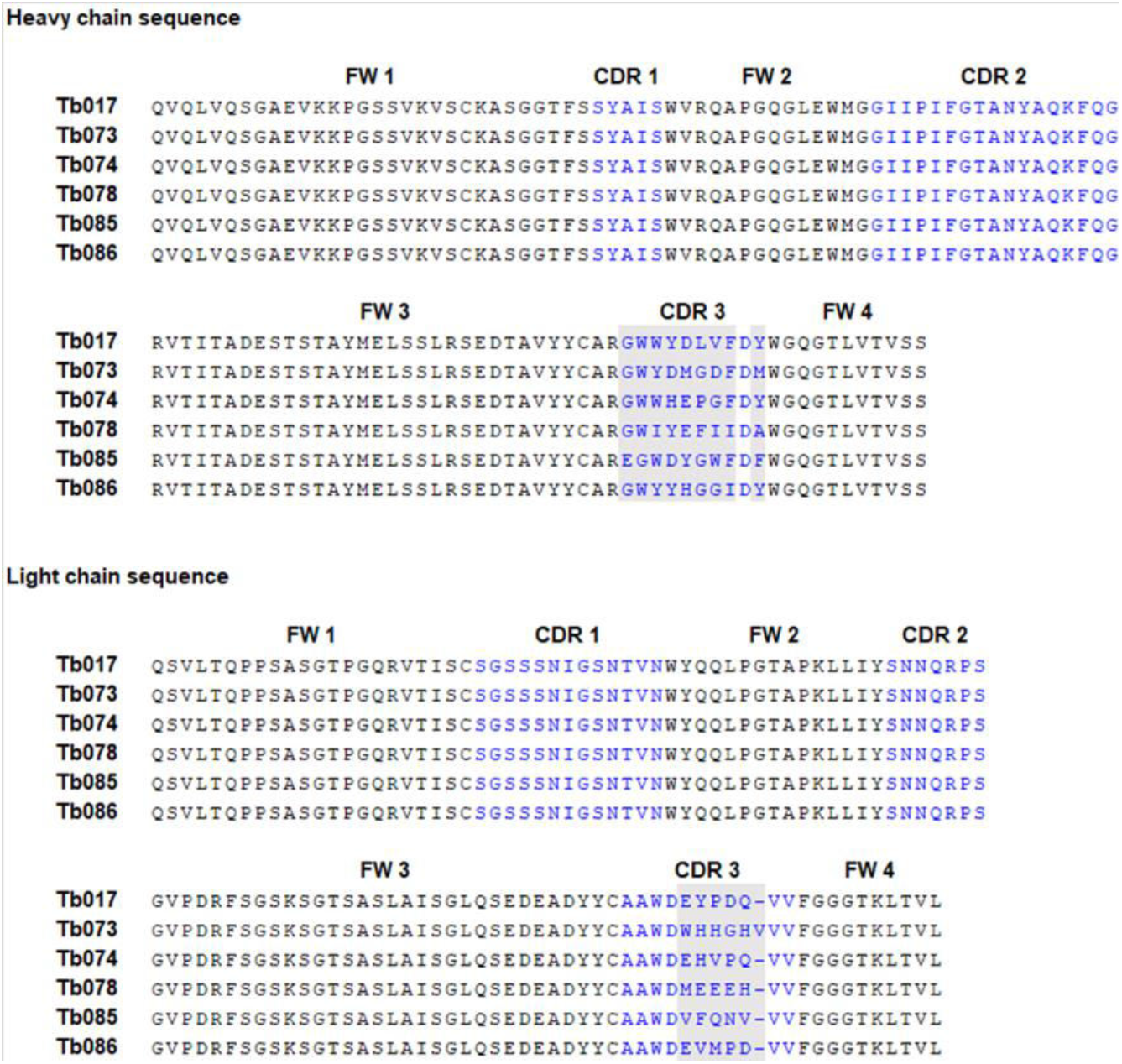
Sequences of the six scFv targeting the HpHbR NTD. The framework domains (FW) are shown in black and the complementarity-determining regions (CDR1-3) are shown in blue. Sequence variation between scFvs is in CDR3, as annotated by grey boxes.

**S2 Figure:**
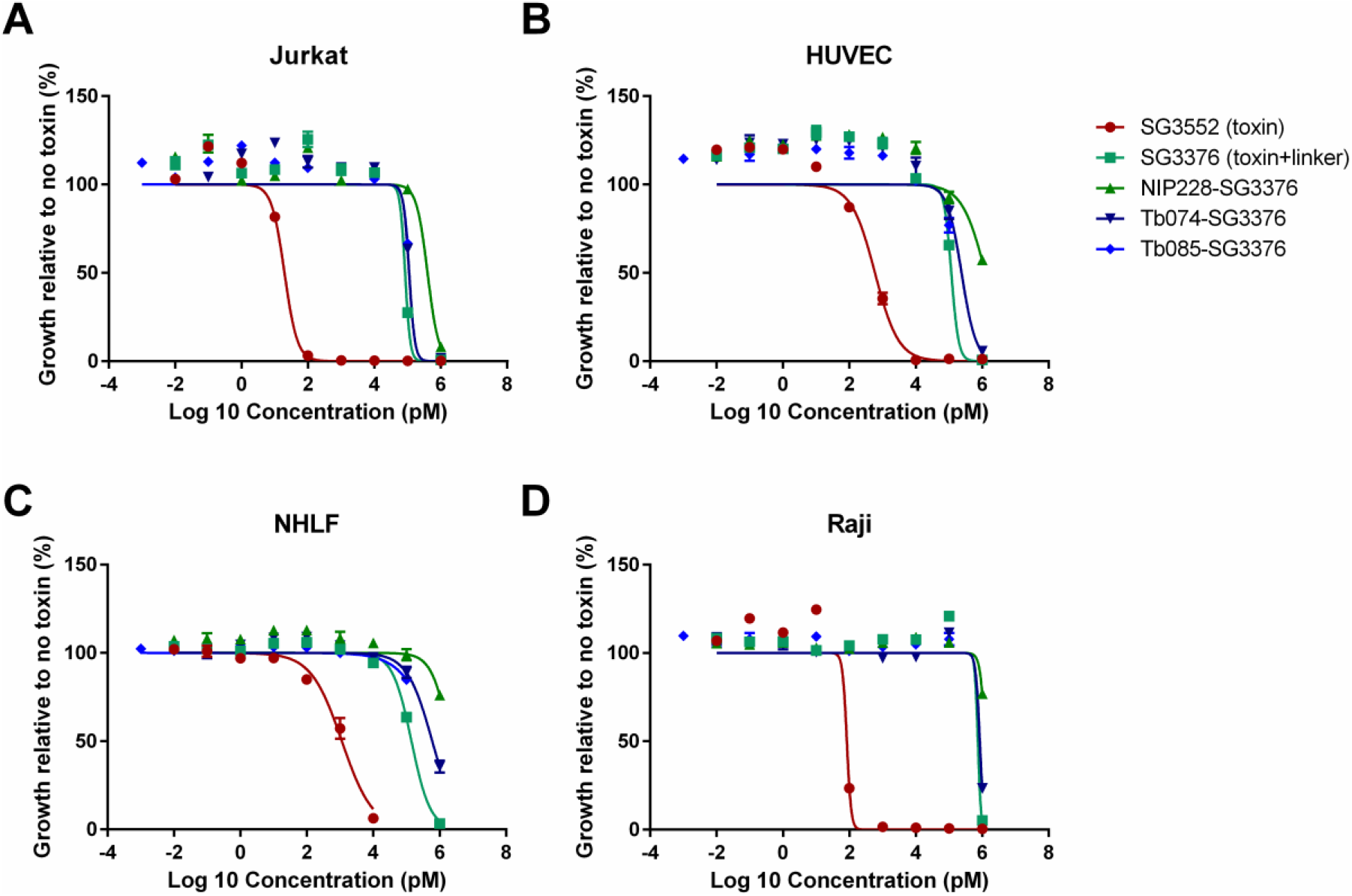
Conjugating toxin SG3552 to antibodies that recognise the *T. brucei* HpHbR reduces toxicity against human cell lines. Toxin SG3552, toxin plus linker SG3376 and the associated ADCs were incubated with (A) Jurkat T-cells, (B) Human Umbilical Vein Endothelial Cells (C) Normal Human Lung Fibroblasts, and (D) Raji B-cell lymphoma cells in FCS. Toxin SG3552 kills the human cell lines at picomolar concentrations. Killing activity is reduced in all cell lines by the addition of the linker (SG3376) or incorporation into a control or Anti-HpHbR ADC (NIP22-SG3376, Tb074-SG3376, Tb085-SG3376) to mid-to-high nanomolar concentrations. Lines represents nonlinear regression lines of best fit on Log_10_ transformed data, although it was not possible to fit accurate lines or calculate IC_50_ values for the ADCs due to lack of saturation of the cell killing assay. All assays were carried out in triplicate over 96 hours. Error bars represent s.e.m., n=3.

**S3 Figure:**
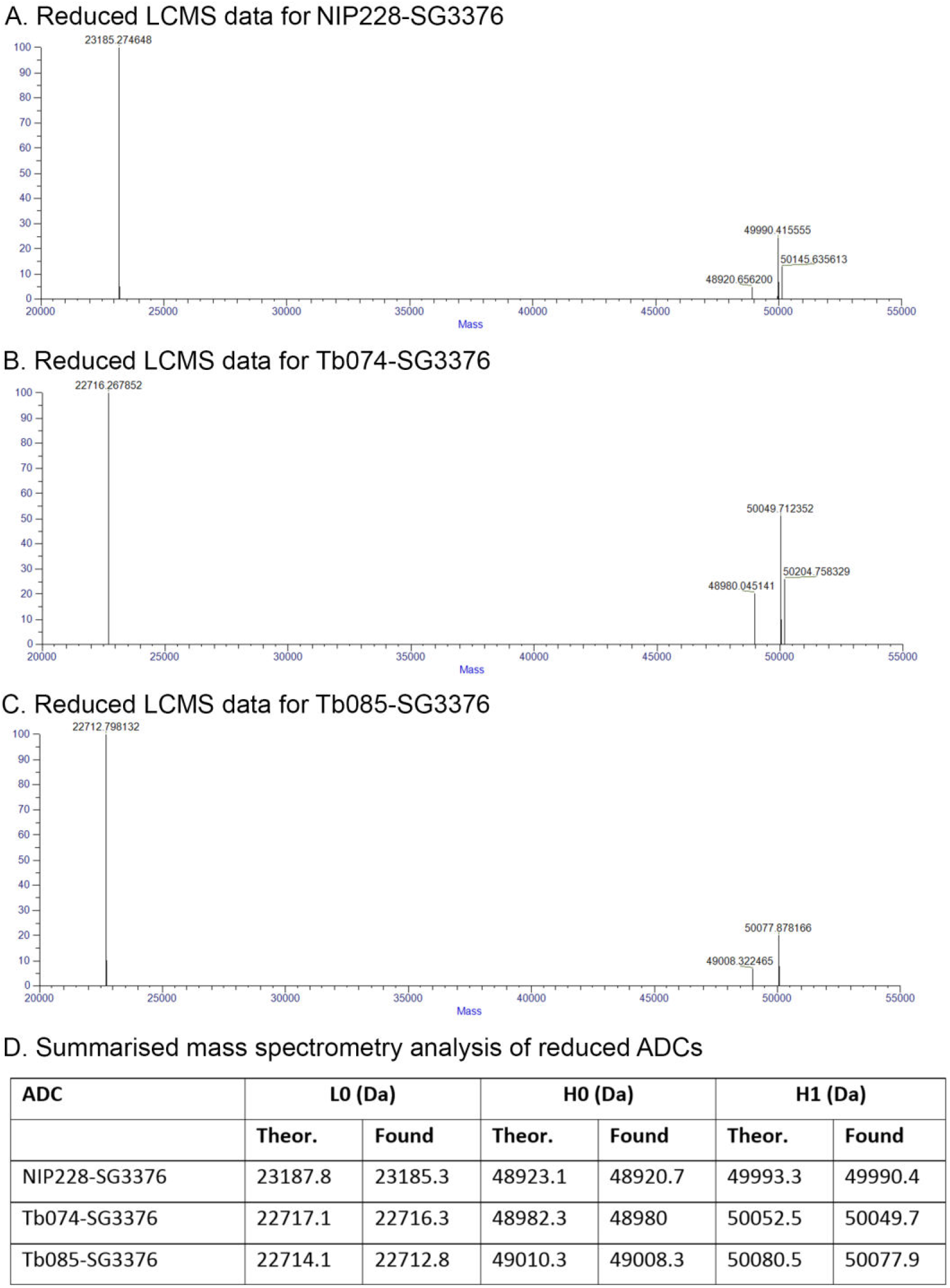
Mass Spectrometry analysis of SG3376-containing antibody-toxin conjugates. Mass spectrometry analysis of reduced antibody-toxin conjugates was performed using a RSLC UPLC system coupled to an Exactive EMR Orbitrap MS. L0 = unconjugated light chain species, H0 = unconjugated heavy chain species, H1 = conjugated heavy chain species.

**S4 Figure:**
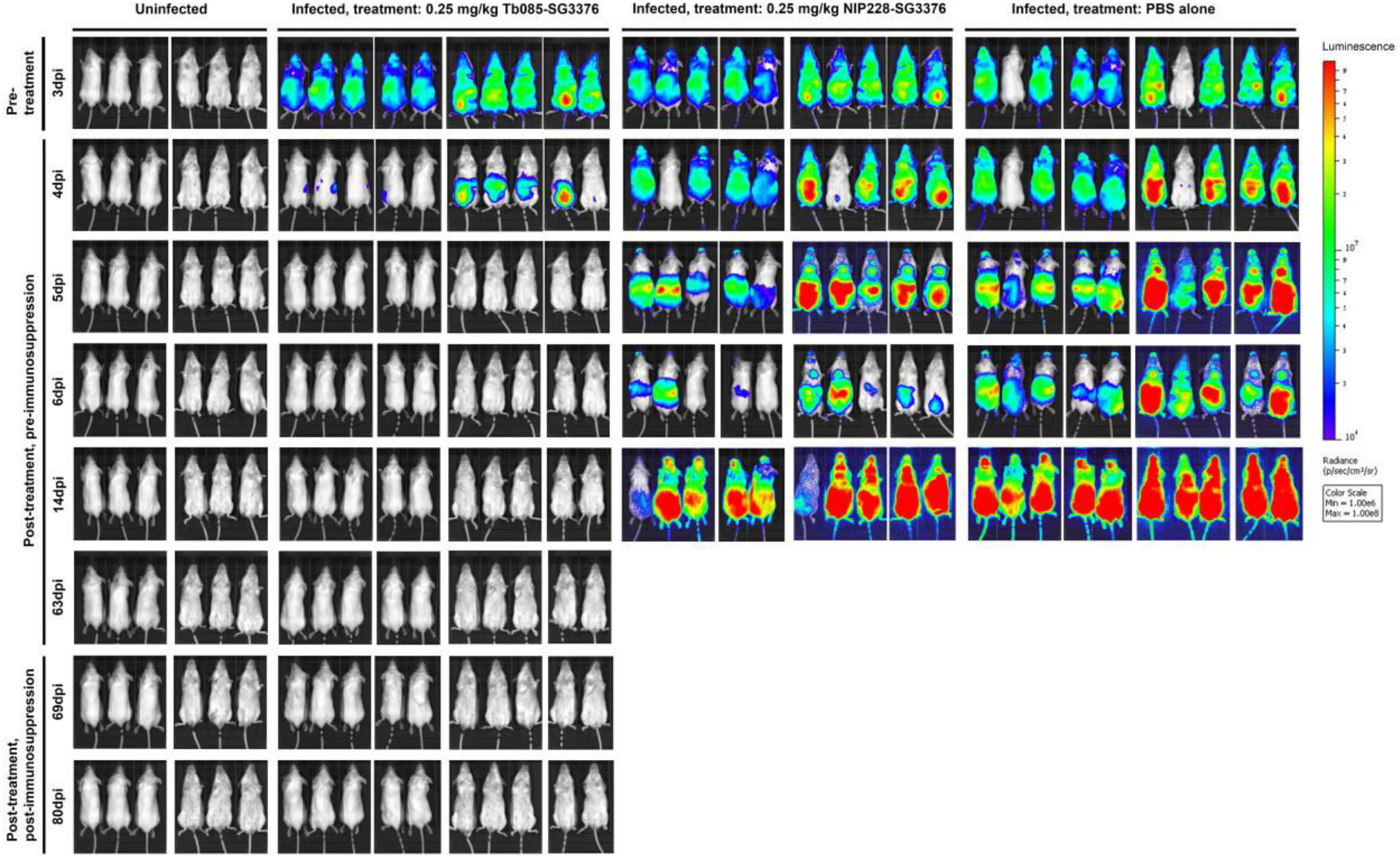
Bioluminescent imaging of *T. b. brucei* infected mice before and after treatment with antibody-toxin conjugates. Parasite burden in mice infected with pleomorphic *T. b. brucei* GVR35-VSL2 cells was assessed by BLI following intraperitoneal injection of d-luciferin. BLI was performed prior to any treatment at 3 days post infection (dpi) and then at regular time points following treatment on 3 dpi with (1) Tb085-SG3376 (n=5), (2) NIP228-SG3376 (n=5) or (3) PBS alone (n=5), with selected time points shown here. Uninfected mice were imaged as controls (n=3). Treatment with Tb085-SG3376 decreased the luminescent signal to that obtained from uninfected control animals within 2 days and this remained the case for the duration of the infection, including following the immunosuppression of Tb085-SG3376-treated mice at 66 dpi. For each group of mice both the dorsal and ventral images are shown. Scale bar represents the photons emitted at any given point on the image. Exposure times range from 0.5 seconds (for heavily burdened mice) to 5 minutes (for uninfected animals). One mouse in the PBS control group had a lower BLI signal than all other infected mice at 3 dpi (S5 Figure). In the image shown here this mouse appears negative, however, this is due to the low exposure time required for adjacent mice. Quantification of the total luminescence from each mouse was also carried out (Figure 4 and S5 Figure).

**S5 Figure:**
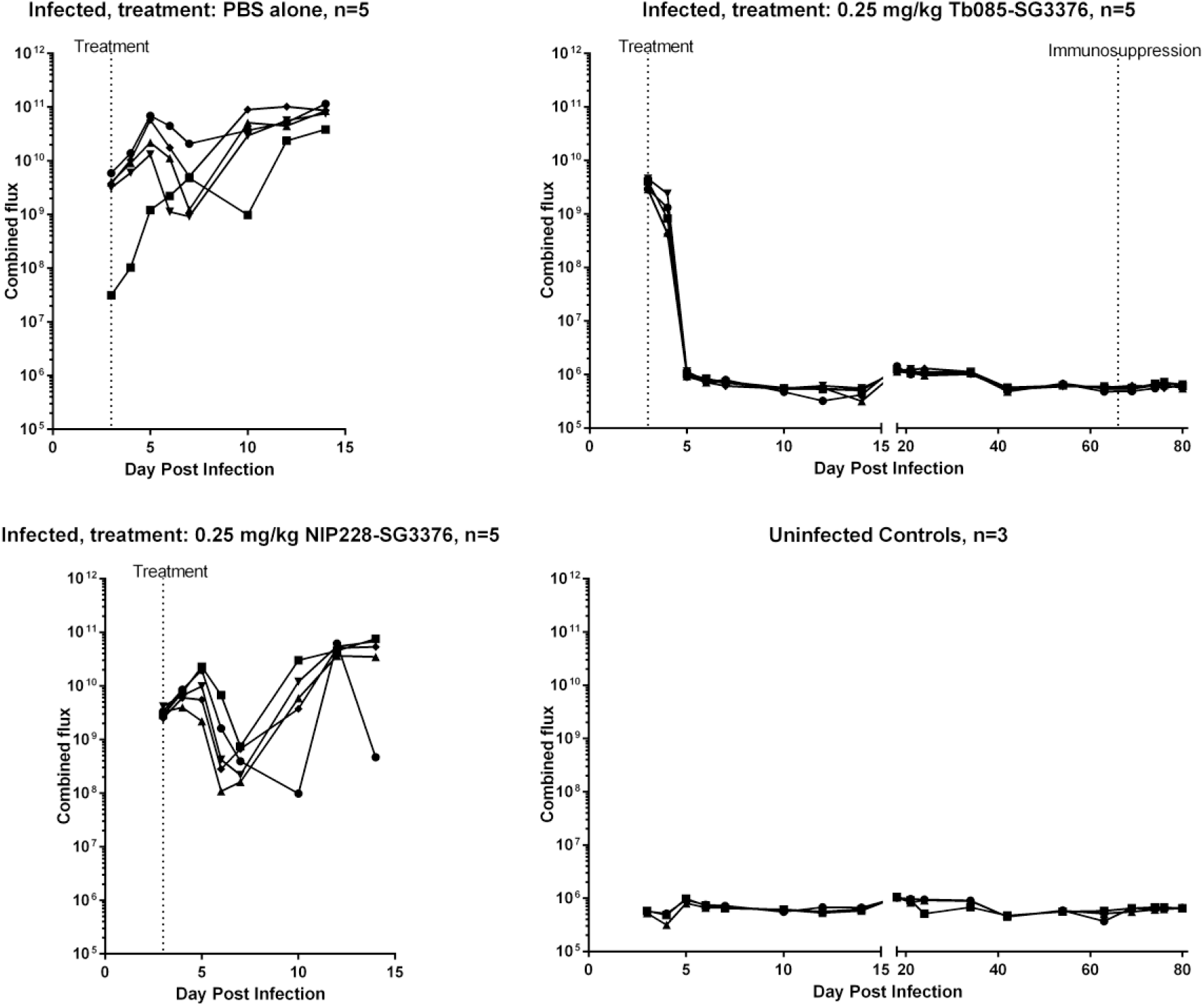
A single low dose of Tb085-SG3376 was able to cure infection in a mouse model of trypanosomiasis: data from individual mice. Three groups of 5 mice were infected with pleomorphic *T. b. brucei* GVR35-VSL2 cells, which allow for parasite burden in live mice to be assessed over a time course by bioluminescent imaging (BLI). BLI was performed prior to any treatment at 3 dpi and then at regular time points following treatment on 3dpi with a single intravenous dose of (1) 0.25 mg/kg Tb085-SG3376 (n=5), (2) 0.25 mg/kg NIP228-SG3376 (n=5) or (3) PBS alone (n=5). Unlike the control-treated mice, Tb085-SG3376 treatment caused a decrease in the luminescent signal to that obtained from uninfected control animals within 2 days and this remained the case for the duration of the infection, including following the immunosuppression of Tb085-SG3376 treated mice at 66 dpi. Mice treated with NIP228-SG3376 or PBS were culled at a humane endpoint on day 14. Quantification shown is the combined (dorsal + ventral) luminescence over the whole mouse in photos per second (p/s). The combined quantification data from the 4 groups of mice are shown in Figure 4. Selected images for the BLI are shown in S5 Figure.

**S6 Figure:**
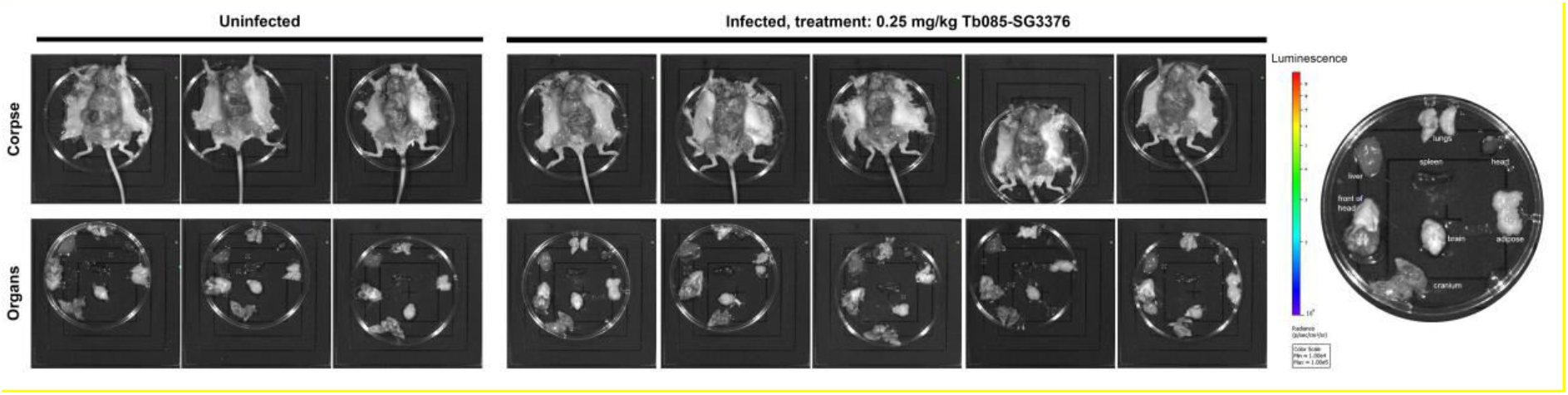
No parasites were detected by BLI post-necropsy in *T. b. brucei* infected mice following treatment with Tb085-SG3376. The five mice that were infected with pleomorphic *T. b. brucei* GVR35-VSL2 cells, treated with 0.25 mg/kg Tb085-SG3376 (3 dpi) and immunosuppressed (66 dpi) were culled at 80 dpi. Post-necropsy, mice corpses and selected organs were assessed by BLI. Consistent with BLI data from live mice, BLI signal was equivalent to the uninfected control mice.

**S1 Table:** IC_50_ values (pM) of SG3199-based toxins and antibody-toxin conjugates against wild type *T. b. brucei*. The IC_50_ values of toxin SG3199, toxin plus linker SG3249, a control ADC (NIP228-SG3249) and five anti-trypanosome antibody toxin conjugates targeting the *T. brucei* HpHbR (Tb017-SG3249, Tb073-SG3249, Tb074-SG3249, Tb078-SG3249, Tb085-SG3249) were calculated against *T. b brucei* wild type (Figure 3). Values in bold are best-fit IC_50_ values, the range is the 95% confidence intervals. All values are shown to 3 significant figures.

| IC50 (pM) |  |
| --- | --- |
|  | *T. b. brucei* |
| **SG3199 Toxin** | **0.86**<br>(0.69-1.08) |
| **SG3249 Toxin plus Linker** | **236**<br>(196-284) |
| **NIP228-SG3249 Control** | **2100**<br>(1760-2500) |
| **Tb017-SG3249** | **32.9**<br>(26.8-40.3) |
| **Tb073-SG3249** | **85.7**<br>(74.6-98.6) |
| **Tb074-SG3249** | **17.3**<br>(14.3-21.1) |
| **Tb078-SG3249** | **74.2**<br>(64.3-85.7) |
| **Tb085-SG3249** | **9.35**<br>(7.59-11.5) |

**S2 Table:** Monomer content and drug-antibody-ratio (DAR) of SG3376-containing Antibody-toxin conjugates. Monomeric purity was determined by size exclusion chromatography (SEC) and the DAR was determined by RP-HPLC. Both assays were performed on a Shimadzu Nexera UPLC system fitted with a Shimadzu Prominence DAD detector. Data were processed using LabSolutions software.

| Antibody toxin conjugate | SEC |  |  | DAR |
| --- | --- | --- | --- | --- |
|  | % HMW | % Monomer | % LMW |  |
| NIP228-SG3376 | 3.2 | 90.1 | 6.7 | 1.77 |
| Tb074-SG3376 | 5.4 | 92.1 | 2.5 | 1.80 |
| Tb085-SG3376 | 0 | 100 | 0 | 1.79 |

